# Polychaete capture by native jellyfish and invasive ctenophore reveals a novel benthic–pelagic trophic link

**DOI:** 10.1101/2025.10.18.683197

**Authors:** Hannah HJ Yeo, Laura Ferreira, Erik Kristensen, A Garm, Jamileh Javidpour

## Abstract

Shallow coastal and estuarine habitats are among the most productive ecosystems, sustained by dynamic benthic–pelagic coupling. While traditionally described through detrital fluxes, living-mediated trophic interactions remain underexplored. Here we present the first field evidence that benthic nereidid polychaetes are preyed upon by gelatinous zooplankton; the native scyphomedusae *Aurelia aurita* and the invasive ctenophore *Mnemiopsis leidyi*. These events occurred most frequently in summer, particularly in the inner reaches of a Danish fjord, and were more often associated with *A. aurita* than *M. leidyi*. Stable isotope analyses revealed seasonal and species-specific patterns: in summer, polychaetes shared similar isotopic signatures with their gelatinous predators, while in autumn they exhibited significant enrichment, possibly reflecting starvation or decay. Mixing models indicated that polychaetes constitute the second most important dietary component for *A. aurita* and *M. leidyi* during summer, after seston and zooplankton, respectively but declined in importance in autumn. These finding uncover a previously overlooked trophic pathway through which benthic prey subsidize pelagic consumers, strengthening benthic–pelagic coupling. Despite their lower observational frequency in *M. leidyi*, dietary models suggest ctenophores may exploit this resource opportunistically, with potential implications for competitive interactions and invasion dynamics. Our results highlight the need to incorporate living benthic–pelagic interactions into ecological models of energy flow and invasion ecology.

## Introduction

Benthic–pelagic coupling is increasingly recognized as a dynamic, bi-directional process, rather than a simple downward detrital flux, that links the seafloor and the water column. Classic studies emphasize that the role of marine snow, faecal pellets, and carcases in sustaining benthic communities (Smith et al. 1989, Smith 1992, Fujioka et al. 1993). Yet, mounting evidence shows that living organisms actively mediate energy and material exchange via their movements, life□history strategies, and feeding behaviour. Gobies, for example, regularly ascend from the benthos to consume pelagic prey such as jellyfish, returning this energy to benthic foodwebs (Utne-Palm et al. 2010). Likewise, crustaceans and other taxa perform diel or seasonal vertical migrations (Raffaelli et al. 2003, Baustian et al. 2014), while physical forces such as storms or tides can resuspend benthic fauna into the pelagic zone (Essink et al. 1989, Garstecki and Wickham 2001). These mechanisms reframe benthic–pelagic coupling as an active living network, rather than a one-way debridal subsidy.

Among the organisms facilitating this coupling, gelatinous zooplankton play a prominent but underappreciated role. Most studies has focused on their functions within the pelagic domain: preying on zooplankton, fish eggs, and larvae (Mistri et al. 2025), and serving as prey for higher□level consumers like turtles, seabirds, and fish (de Lafontaine and Leggett 1988, Arai 2005, Dodge et al. 2011, Jarman et al. 2013). Medusae and other gelatinous taxa also host a remarkable diversity of associates, including fishes and invertebrates, engaging in parasitism, commensalism, mutualism, and kleptoparasites (Fleming et al. 2014, Riascos et al. 2015, Ohtsuka et al. 2009, Ingram et al. 2017). Juvenile fish (Masuda et al. 2008; D’Ambra et al. 2015) and crabs (Sal Moyano et al. 2012) often use medusae for protection or nurseries, while others consume their tissue or captured prey (Miyajima et al. 2011, O’Rorke et al. 2015, Towanda and Thuesen 2006, Weil et al. 2019). Some amphipods even deposit eggs within medusae, with their young feeding on host tissue or stolen prey (Gasca and Haddock 2004, Harbison et al. 1977, Dittrich 1988). Others, including ophiuroids and crustaceans, use medusae for transport to new areas (Marliave and Mills 1993, Kanagaraj et al. 2008, Sal Moyano et al. 2012). By serving simultaneously as food source, habitat, nursery, and transport vector, medusae function as mobile ecological hubs that connect benthic and pelagic realms (Doyle et al. 2014). Despite the diversity of known medusa–associate interactions, trophic links to benthic organisms remain largely undocumented. While associations with fishes and crustaceans have received attention, the roles of sediment□dwelling invertebrates such as adult polychaetes are rarely explored. These worms are almost absent from medusa–associate literature (but see polychaete larvae in Table 4 of Mutlu 2001), despite their high abundance, trophic versatility, and foundational role in benthic food webs. Investigating such interactions may reveal overlooked pathways through which gelatinous zooplankton engage in benthic–pelagic coupling.

Here, we document nereidid polychaetes (e.g. *Platynereis dumerilii* and *Alitta succinea*) pray upon by two gelatinous zooplankton: the native scyphozoan *Aurelia aurita*, and the invasive ctenophore *Mnemiopsis leidyi*. These taxa differ markedly in their feeding strategies. *A. aurita* is a seasonal, moderate□intensity predator, while *M. leidyi* is a voracious, continuous feeder capable of restructuring pelagic communities (Piccardi et al. 2025). To our knowledge, this is the first report of adult Nereididae being consumed by either cnidarian medusae or ctenophores. This finding prompts several questions: Are nereidid polychaetes opportunistically preyed upon at the seabed, or are they captured during their heteronereis-stage surface swarming, coinciding with seasonal gelatinous blooms? What is their dietary contribution to these gelatinous predators? To address these questions, we (i) documented nereid polychaetes (family Nereididae) capture by *A. aurita* and *M. leidyi*, ii) evaluated host-specific and seasonal patterns of predation, and (iii) used stable isotope analysis and dietary mixing models to assess the trophic contribution of polychaetes. This study provides insights into benthic-pelagic energy pathways and highlights the potential for invasive species to reshape trophic interactions via novel resource use.

## Materials and methods

### Study system

This study was conducted in the inner Danish estuarine complex of Kerteminde Fjord/Kertinge Nor, which connects to the Great Belt via a narrow channel (Figure 1). The system exhibits pronounced seasonal variability in temperature (0–25□°C) and salinity (10–25 psu) and is classified as moderately eutrophic due to sustained nutrient inputs from surrounding agriculture and urban areas (Riisgård et al. 1995). Bloom-forming gelatinous zooplankton are a prominent seasonal feature, with *A. aurita* typically present from May through September and *M. leidyi* from July to October (Riisgård et al. 2010).

**Figure 1.**
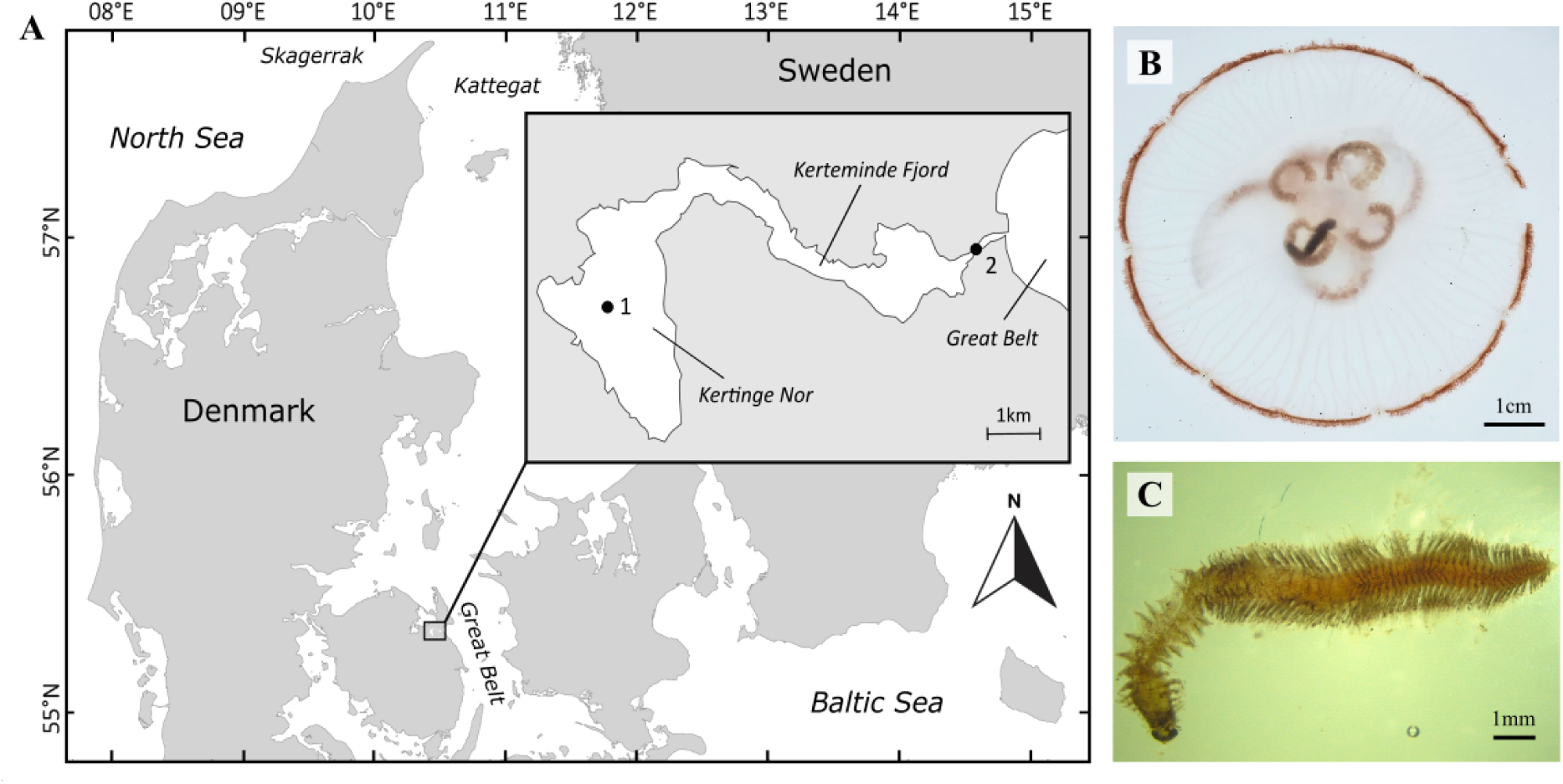
(A) Location of the two sampling sites surveyed during the 2022 fieldwork. (B) Photograph showing the association of *Aurelia aurita* with a Nereid polychaete. (C) Microscopic view of the same polychaete shown in panel B, identified as *Platynereis dumerilii*.

### Field sampling of gelatinous zooplankton and polychaetes

*A. aurita*, *M. leidyi* and free-living nereidid polychaetes were collected near the water surface using a handheld dip net from a small boat at two locations: Site1 in the inner Kertinge Nor, and Site2 near the mouth of Kerteminde Fjord (Figure 1A). Sampling took place from June to November 2022 and targeted a minimum of five individuals of each species per session to enable seasonal comparisons. Collection points were randomized within each site, with the goal of obtaining individuals both with and without captured Nereididae. Immediately after sampling, specimens were rinsed thoroughly with filtered seawater (0.2µm) to remove debris and epibionts, measured, and dissected. Bell (mesoglea) tissue was excised and dried at 60°C for 48 h. Any nereidid polychaetes observed inside the gelatinous tissues were carefully removed, recorded, and processed using identical drying and storage protocols. Site 1 was revisited in summer 2025 to confirm the identification of captured polychaetes in 12 new samples (Figure 1B – C); these were not processed for stable isotope analysis.

### Zooplankton sampling and processing

Plankton were sampled with vertical tows of 20 µm and 100 µm mesh nylon nets, each equipped with a 35 mL cod-end cylinder with a valve. Nets were lowered to a depth of 2 m for 30 s and retrieved at a constant speed. Upon retrieval, nets were rinsed with filtered seawater and the samples transferred into labeled containers kept on ice during transport. In the laboratory, samples were sieved through appropriate mesh sizes (100 µm and 200 µm, respectively) to remove detritus. Non gelatinous zooplankton (hereafter referred to as zooplankton) were chilled at –20°C for 30–60 min to immobilize them before sorting under a stereo microscope. For each taxon, 50 individuals were pooled as one composite sample, placed into prelabeled Eppendorf tubes, and oven dried at 60°C for 48 h.

### Seston sampling and processing

Seston was sampled in situ with a 5 L Niskin bottle at 2 m depth at each site and time point. Samples were kept on ice during transport. In the laboratory, 2 L of seawater per sample was filtered using a glass vacuum filtration system onto pre-combusted GF/F filters (450°C for 3 h). Filtration units and filters were rinsed with 10% HCl between samples and subsequently neutralized with deionized water following Howe and Simenstad (2011). Filters were folded, sealed in sterile petri dishes, stored at –80 °C, and later oven dried at 60 °C for 48 h.

### Sample preparation for isotope analysis

All dried tissues were homogenized to a fine powder with an agate mortar and pestle. For gelatinous zooplankton and polychaetes, 1–6 mg (±0.2 mg) of dry material was encapsulated in tin capsules. For seston, four subsamples were cut from each filter using a sterile isotopic cutter and encapsulated. Capsules were analyzed for δ¹³C and δ¹□N using isotope ratio mass spectrometry (IRMS) at the University of Southern Denmark. Total biomass on filters was calculated from known filter areas and subsample volumes.

### Stable isotope data processing and statistical analyses

All isotope data were preprocessed by removing records lacking δ¹³C, δ¹□N, or C:N ratio information. Outliers with values deviating by an order of magnitude, likely due to contamination or analytical error, were excluded. To account for lipid effects, C:N ratios were evaluated against a 3.5 threshold, and δ¹³C values were lipid normalized where necessary following Kiljunen et al. (2006). Relationships between gelatinous zooplankton size and Nereididae counts were visualized with histograms and density plots in RStudio. Isotope data were analyzed using generalized linear models (GLMs) with a quasi-likelihood approach to account for violations of normality and heteroscedasticity. Separate models were fitted for δ¹³C and δ¹□N, with Site (categorical, two levels) and Season (categorical, two levels) as predictors. Model selection and simplification were carried out iteratively until all remaining interaction terms in the model were statistically significant. Pairwise comparisons were conducted using the *emmeans* package (Lenth 2025) and corrected for multiple testing using the *Tukey* adjustment.

### Trophic discrimination factors (TEFs) and trophic position

Few studies report TEFs for gelatinous zooplankton. We considered values from D’Ambra et al. (2014), Tilves et al. (2018), and Schaub et al. (2021) and selected those representing jellyfish feeding on crustacean zooplankton (Schaub et al. 2021: Δδ¹³C = 1.19 ‰; Δδ¹□N = 2.09 ‰). However, because the *tRophicPosition* package in R only implements TEFs from Post (2002) and McCutchan et al. (2003), we applied the closest available values: Δδ¹³C = 1.3 ± 0.3 ‰ (muscle tissue) and Δδ¹□N = 2.1 ± 0.21 ‰ (whole tissue) (McCutchan et al. 2003). Trophic positions were modeled using the “oneBaseline” Bayesian model in *tRophicPosition* (Quezada-Romegialli et al. 2018) with 20,000 adaptive samples, 20,000 iterations, 2,000 burn-ins, and five chains.

### Dietary mixing models

To estimate the proportional dietary contributions to *A. aurita*, we implemented Bayesian mixing models in *MixSIAR* (v3.1; Stock and Semmens 2016). Nereididae, seston, and zooplankton were from isotopic biplots identified as the most probable food sources and incorporated into the mixing models. Uninformative priors were used with 1000 chain length, 500 burn-ins, three chains, and trophic enrichment factors from Schaub et al. (2021) were applied. Model convergence and posterior distributions were evaluated following MixSIAR guidelines.

## Results

### Field observations

From June to October of 2020 and 2021, we documented 68 separate instances of nereidid polychaetes entangled in the oral arms or bells of gelatinous zooplankton. Out of 152 *A. aurita* individuals examined, 38 (25%) contained at least one nereidid worm (Table S1). In contrast, only 3 of 67 *M. leidyi* (4%) individuals contained nereidids. Polychaetes were typically located beneath the bell, embedded within the mesoglea, or inside the gastric cavity of *A. aurita*. Most captured worms appeared intact (Fig. 1), although some showed signs of partial digestion or degradation. *A. aurita* individuals hosting nereidids were slightly smaller on average than those without (Figure S1), though no size pattern was evident for *M. leidyi*, potentially due to limited sample size (Figure S2). The maximum number of nereidids per medusa was two, observed in *A. aurita* individuals with bell diameters between 52 and 84□mm (Figure S4). Additionally, nereidids recovered from jellyfish were marginally larger than free-living individuals collected during the same period (Figure S3). Among all identified worms, 91% were *Platynereis dumerilii* and 9% were *Alitta succinea*.

### Trophic relationships between nereidid polychaetes and gelatinous predators

Isotopic biplots and generalized linear models (GLMs) revealed significant seasonal variation in trophic patterns between nereidid worms and gelatinous zooplankton (*p* =2.35e^-14^). During summer, the isotopic niches of Nereididae and *A. aurita* overlapped broadly, suggesting a trophic link or shared resource use. In contrast, this overlap disappeared in autumn (Figure 2), reflecting a shift in resource use or metabolic status. At Site 1 in summer, free-living Nereididae and those found inside *A. aurita* exhibited δ¹³C values closely matching those of the jellyfish host, with an average enrichment of ∼4–8.7‰ relative to seston and zooplankton (*p*□<□0.05; Table S3). The δ¹□N values of Nereididae inside A. aurita were ∼0.35‰ higher than free-living conspecifics, though this difference was not statistically significant (Table S3). Importantly, the δ¹□N signatures of these worms closely mirrored those of their A. aurita hosts (Figure 2A), supporting a tight trophic coupling during summer. Across all seasons and sites, *M. leidyi* consistently showed higher δ¹□N enrichment and trophic position than *A. aurita,* with the exception of Site 2 in autumn, where no *A. aurita* individuals were present (Figure 2,3). In autumn, isotopic pattern shifted: Nereididae inside *A. aurita* were significantly enriched in δ¹³C (by 4.0–4.3‰, *p* = 0.002; Table S4) and also showed elevated δ¹□N values (by 1.8–2.2‰), although the later was not statistically significant (*p* = 0.168; Table S4). These enrichments correspond to an apparent trophic offset of nearly one full level above their jellyfish hosts (Figure 3), suggesting either metabolic fractionation or degradation effects. Additionally, both A. aurita and M. leidyi at Site 1 in autumn were less enriched in δ¹³C compared to their summer counterparts, indicating seasonal shifts in carbon source utilization.

**Figure 2.**
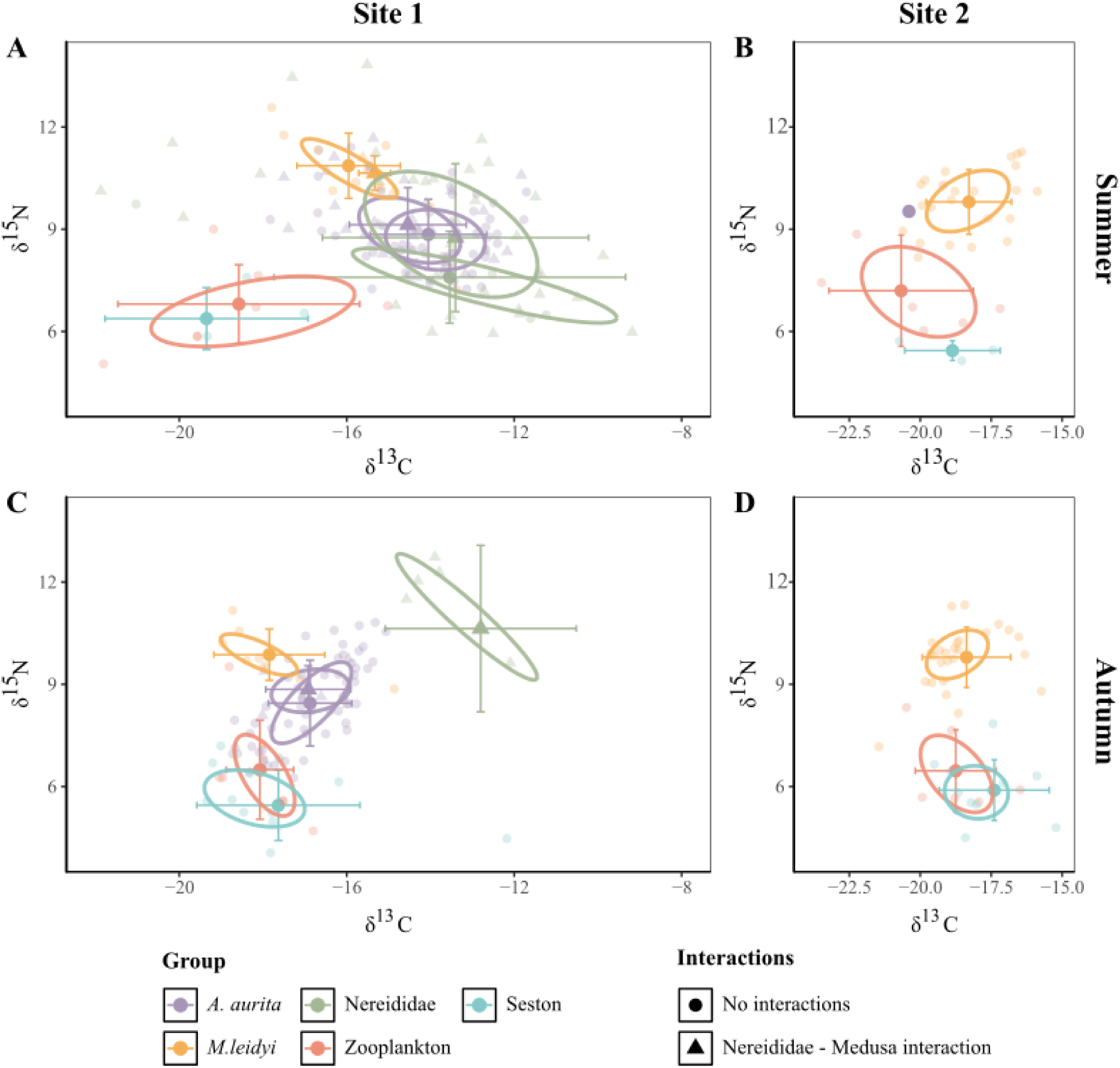
Biplots of d13C and d15N isotope values with 40% ellipse areas (representing the standard isotope niche) during A) Summer at Site 1, B) Summer at Site 2, C) Autum at Site 1 and D) Autumn at Site 2.

**Figure 3.**
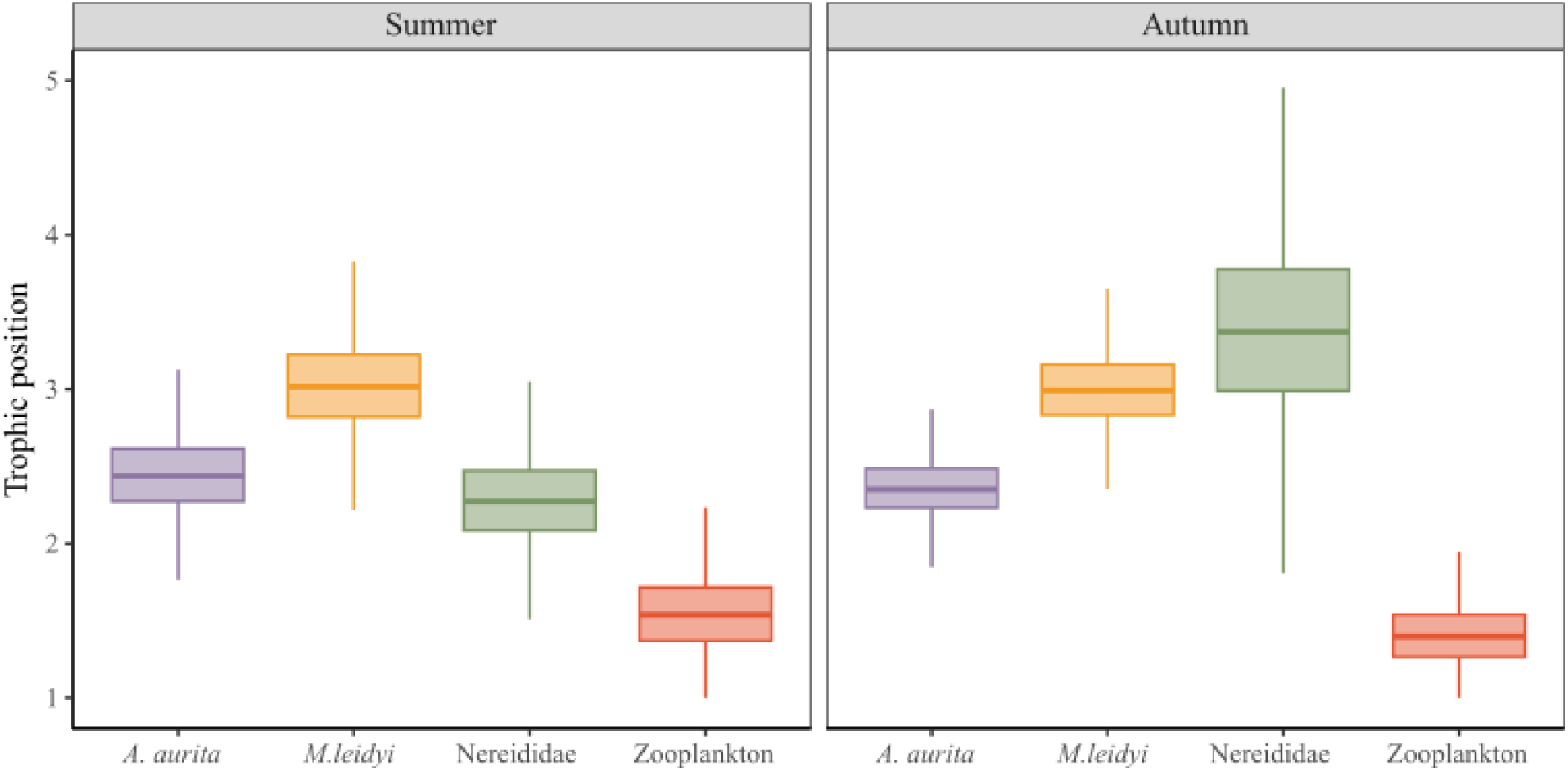
Variation in bayesian estimated trophic position of gelatinous zooplankton species, non gelatinous zooplankton and Nerridae during the Summer and Autumn.

### Dietary contributions

Bayesian MixSIAR models revealed distinct seasonal and species-specific diatery pattern (Figure 4). During summer, seston represented the dominant dietary source of *A. aurita*, followed by Nereididae polycheates and to a lesser extent, zooplankton (Figure□4A). In contrast, *M. leidyi* derived a substantial portion of its assimilated diet from zooplankton with Nereididae contributing secondarily and seston playing a minor role (Figure□4B). This patern shifted markedly in autumn, when both *A. aurita* and *M. leidyi* exhibited diet dominated almost exclusively by zooplankton, consistent with a reversion to typical pelagic feeding (Figures 4C, 4D). The near absence of seston and nereidid contributions in autumn models corresponds to reduced benthic-pelagic trophic coupling during this period.

**Figure 4.**
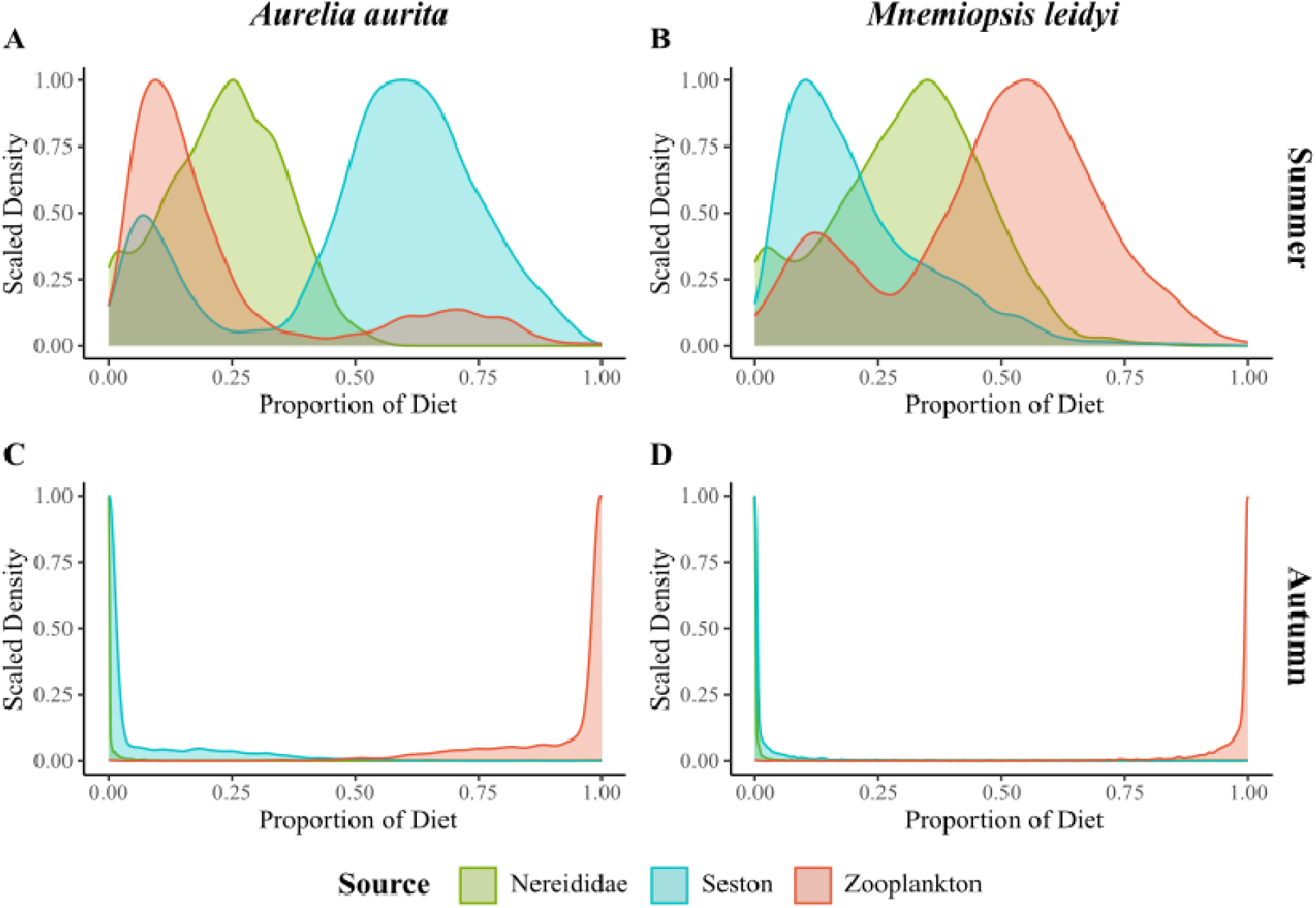
Bayesian estimated proportional dietary contributions for *A. aurita* and *M*. *leidyi* individuals at Site 1, generated using MixSIAR. The dietary model includes the Nereididae polychaete, seston and zooplankton. (A) proportion of diet for *A. aurita* during summer; (B) proportion of diet for *M*. *leidyi* during summer; (C) proportion of diet for *A. aurita* during autumn; and (D) proportion of diet for *M*. *leidyi* during autumn.

## Discussion

Here we present the first evidence of nereidid polychaetes captured by the native scyphozoan jellyfish *Aurelia aurita* and, to a lesser extent (i.e. based on fewer observations), with the invasive ctenophore *Mnemiopsis leidyi*. *A. aurita*, a widely distributed and extensively studied genus of scyphozoan jellyfish, has not previously been observed with nereidid worms in the gut. This lack of prior documentation may indicate that the incidence is rare or recently emerged due to shifts in environmental or ecological dynamics, for example, changes that align predator blooms and benthic prey spawning cycles. Alternatively, the interaction may have been previously overlooked because of the short digestion time of prey in *A. aurita*; our preliminary observations suggest that Nereididae polychaetes are digested in approximately two hours, or possibly less (pers. observation). This new record of pelagic-stage gelatinous zooplankton feeding on benthic polychaetes highlights a previously overlooked pathway in benthic–pelagic coupling. By consuming polychaetes, jellyfish and ctenophores effectively redirect benthic-derived energy – energy that would otherwise be lost to the seafloor following polychaete mortality – back into the water column.

### Polychaete identification and capture mechanisms by gelatinous zooplankton

Although the polychaetes collected in 2022 were not identified to species, their relatively small body sizes (8–30 mm) offered a useful clue. Three nereidid species are commonly found in the shallow soft sediments of the Kerteminde Fjord/Kertinge Nor system—*Alitta virens*, *Hediste diversicolor*, and *Alitta succinea* (Miron and Kristensen 1993). Since adult individuals in the benthic stage of all three species typically reach much larger sizes (75–130 mm; Kristensen 1984), their presence in our samples is highly unlikely. This interpretation is further supported by re-sampling and species-level identification conducted in 2025, which confirmed the presence of small spawning heteronereis individuals of *A. succinea* and *Platynereis dumerilii* captured by *A. aurita* at the same study sites.

We believe that these benthic-pelagic linkages happen through one primary mechanism. Gelatinous zooplankton capture and ingest polychaetes as prey within the water column or near the water surface when they leave their burrows or tubes to reproduce. It has been recorded that metamorphosed sexual heteronereis individuals of both *A. succinea* and *P. dumerilii* mature individuals will leave the benthos and swim to the surface to spawn. This spawning is often influence by the lunar cycle and happens more frequently during spring and summer (Korringa 1947, Grant 1989, Ram et al. 2008, Özpolat et al. 2021), following which, the adults die. This timing coincides with the period when Nereidid individuals were observed most frequently inside the gelatinous zooplankton (Table S2). Furthermore, we found that most polychaetes inside *A. aurita* that were heteronereis stages of *P. dumerilii*; for instance, enlarged adult eyes and change in colour (Fischer et al. 2010, Vervoort and Gazave 2022), and we also noted the presence of gametes in some individuals.

### Trophic dynamics from stable isotopes – dietary proportions and seasonal variation

Stable isotope analysis revealed that during summer, Nereididae, were less enriched in ¹□N and occupied a lower trophic position compared to *A. aurita* and *M. leidyi*. The similarity in ¹³C signatures between Nereididae and *A. aurita* suggests that the medusa likely fed on these polychaetes during this period. This interpretation was further supported by mixing models, which quantified their relative dietary contributions. An alternative explanation for the elevated δ¹□N values observed in *A. aurita* and particularly in *M. leidyi* could be the ingestion of fish eggs or larvae, which are typically abundant in temperate estuarine systems during summer and early autumn. Seasonal surveys in Scandinavian and North Sea fjords show that larval fish densities peak from late spring through mid-summer, with retention of larvae in inner fjord basins where both gelatinous species were collected (e.g. Pacariz et al. 2014, Energinet 2024). Occasional consumption of such high-trophic prey would transiently increase δ¹□N enrichment in gelatinous zooplankton, even without major dietary shifts. However, this explanation is less likely to account for the consistent δ¹□N overlap between *A. aurita* and nereidids in summer and the limited number of large larval prey observed in gut inspections. Nonetheless, we cannot rule out sporadic ingestion of larval fish as a contributing factor, particularly for *M. leidyi*, which is known to consume ichthyoplankton under dense larval conditions (Purcell et al. 2001; Javidpour et al. 2009). Future work should therefore include concurrent sampling of fish larvae and isotopic baselines to better resolve this potential trophic link.

Zooplankton, the primary food source of *A. aurita* and *M. leidyi* (Sullivan & Gifford 2004, Javidpour et al. 2016), remained the dominant prey in autumn, consistent with our mixing model outputs. In contrast, during summer, polychaetes represented a substantial portion of the diet for both species, ranking as the second most important food source. For *A. aurita*, the summer diet shifted more strongly toward seston (as also reported by Javidpour et al. 2016), with polychaetes providing a strong complementary contribution. Meanwhile, *M. leidyi* continued to rely primarily on zooplankton in summer, followed by polychaetes and, to a lesser extent, seston.

An interesting result from this study was the enriched ¹□N levels of polychaetas captured by *A. aurita* during Autumn. Since δ¹□N generally increases with trophic level, and thus the polychaetas are also reflected as a trophic level above the medusa predator (see Figure 4). However, given the established feeding ecology and biology of these polychaetes—primarily detritus and microalgae feeders; the Nereids as predator of jellyfish explanation is unlikely (see also dietary studies on other nereid polychaetes, e.g. Copeland and Wieman 1924; Wells and Dales 1951; Tenore and Gopalan 1974; Heip and Herman 1979; Costa et al. 2006). Furthermore, we had observations of polychaetes being digested by the jellyfish, and it is impossible that heteronereis stages of the two nereidid polychaetes feed on living gelatinous zooplankton. The juvenile stage of these benthic worms occupy U-shaped burrows in the sediment or construct mucous tubes near macroalgae or seagrass bed (in the case for *Platynereis dumerilii*). Their feeding ecology is primarily detritivorous and algivorous, as they scavenge the upper sediment layer enriched in detritus, microphytobenthos, and fragments of macroalgae (Gambi et al. 2000, Gillet et al. 2011, Özpolat et al. 2021). Therefore, the enriched δ¹□N levels, leading to a higher trophic level observed in polychaetes must have another explanation, rather than direct feeding on gelatinous zooplankton.

Notably, we observed heteronereis individuals with gametes present within the tissues of *A. aurita*, which leads us to think that the elevated δ¹□N levels may be linked to physiological changes or heightened energy demands during reproductive metamorphosis. In *P. dumerilii*, the onset of sexual metamorphosis is characterized by cessation of feeding, extensive morphological changes, and emergence from the tube to swim to the surface, where spawning occurs before death (Fischer et al. 2010). Stable nitrogen isotope enrichment has previously been associated with catabolic states such as fasting, nutritional stress (reflecting suboptimal health), and processes related to death and decomposition (Hobson et al. 1993, Gannes et al. 1997, Loudon et al. 2007, Kempster et al. 2007, Keenan & DeBruyn 2019). Accordingly, the δ¹□N enrichment observed here could reflect several potential scenarios: (i) starvation stress in still-living individuals, (ii) active spawning with high energetic costs, or (iii) later stages of deterioration associated with death and decomposition. Notably, free-living polychaetes show slightly lower δ¹□N enrichment compared to those ingested by jellyfish that are likely in more advance stages of decay or partial digestion. To disentangle these possibilities, future work should more carefully document the physiological condition and lividity (time since death) of polychaetes prior to dissection and preservation and relate these observations to δ¹□N variation across intermediate stages of the reproductive process.

Another interesting result from this study is the decrease in stable carbon isotope values of both *A. aurita* and *M. leidyi* from summer to autumn inside Kertinge Nor (Site 1). Seasonal variation in δ¹³C values has been recently documented for A. aurita polyps, which exhibit a negative shift in δ¹³C during colder months (Lucas et al. 2025). Although polyps and medusae are distinct generations, such isotopic patterns may nevertheless be reflected in medusae if cohorts originate from different strobilation events that occur under seasonally varying baseline conditions. Medusae released later in the season would thus inherit the isotopic signature of polyps exposed to more depleted carbon sources, such as benthic-derived or terrestrial organic inputs, that dominate in autumn.

This pattern implies that the observed seasonal δ¹³C decline in medusae does not reflect physiological change within individuals but rather cohort replacement driven by temporal variability in polyp carbon sources. Such variability could arise from increased benthic remineralization, enhanced terrestrial inflows, or microbial recycling of organic matter that is typically ¹³C-depleted. Similar pelagic-to-benthic seasonal shifts in carbon sourcing have been reported in other temperate systems (Grey et al. 2001; Javidpour et al. 2016), supporting the idea that isotopic baselines propagate through successive life stages and feeding pathways within estuarine food webs.

### Resource competition and future implications of invasive species

Although field observations and gut content analyses revealed more frequent captures of Nereididae by *A. aurita* (38 samples; 12 observations in 2025) than by *M. leidyi* (3 samples), our stable isotope mixing models indicated a higher proportional contribution of Nereididae to the diet of *M. leidyi* during summer. This apparent discrepancy reflects the different temporal scales captured by the two approaches: direct observations provide snapshots of immediate feeding events, whereas isotopic models integrate assimilated diet over longer periods, representing cumulative resource use. Hence, while *A. aurita* may opportunistically ingest polychaete larvae when encountered, *M. leidyi* appears to rely more consistently on this benthic-derived food source through time. Such a feeding strategy suggests that the invasive ctenophore is capable of exploiting benthic–pelagic energy pathways more efficiently than the native medusa. If these interactions become more frequent, they could establish a supplementary trophic route that supports *M. leidyi* persistence and competitive dominance under changing environmental conditions—potentially to the detriment of *A. aurita* populations. We therefore emphasize the importance of expanding future research to quantify the prevalence, timing, and energetic significance of these interactions. Understanding the magnitude of benthic carbon inputs to gelatinous zooplankton food webs will be critical for predicting how invasive and native species partition resources, and for assessing their resilience or vulnerability within increasingly dynamic coastal ecosystems.

## Significance statement

Oikos provides an ideal journal for this study given its focus on novel mechanisms that shape ecological processes across levels of organization and its readership’s strong interest in cross-system linkages, species interactions, and invasion biology. Our findings build directly on prior work on gelatinous zooplankton feeding ecology (e.g. Javidpour et al. 2016) and benthic polychaetes (e.g. Kristensen 1984, Miron & Kristensen 1993), yet these two components have rarely been documented together. This manuscript is therefore of broad relevance to Oikos readers seeking to understand how overlooked interactions influence energy flow, community structure, and ecosystem resilience in coastal systems.

## Data archiving statement

We confirm that, should the manuscript be accepted, the data supporting the results will be archived in an appropriate public repository and the data DOI will be included at the end of the article.

## Ethics statement

The authors confirm that all fieldwork and specimen collections were conducted in compliance with local regulations and with the necessary approvals and permissions from the relevant authorities at the study sites. No additional permits were required for the use of the organisms studied, and no museum specimens were used in this research.

## Acknowledgements

We thank Ivan Fedutin for assistance with fieldwork, and Janus Tem Jensen for mass spectrometry laboratory processing..

## References

Arai MN (2005) Predation on fish larvae by moon jellyfish *Aurelia aurita* under low dissolved oxygen concentrations. Fish Sci 71:748–753.

Baustian MM, Hansen GJA, de Kluijver A, Robinson K, Henry EN, Knoll LB, Rose KC, Carey CC (2014) Linking the bottom to the top in aquatic ecosystems: mechanisms and stressors of benthic-pelagic coupling. Eco-DAS X Symposium Proceedings 25–47.

Ceh J, Gonzalez J, Pacheco AS, Riascos JM (2015) The elusive life cycle of scyphozoan Gelatinous zooplankton – metagenesis revisited. Sci Rep 5:12037. doi: 10.1038/srep12037

Copeland M, Wieman HL (1924) The chemical sense and feeding behavior of *Nereis virens*. Sars. Biol Bull 47:231–238. doi: 10.2307/1536502

Costa PF, Oliveira RF, Fonseca LC da, Jardim R (2006) Feeding ecology of *Nereis diversicolor* (O.F. Müller) (Annelida, Polychaeta) on estuarine and lagoon environments in the southwest coast of Portugal. Pan-Am J Aquat Sci 104–113.

D’Ambra I, Carmichael RH, Graham WM (2014) Determination of δ13C and δ15N and trophic fractionation in gelatinous zooplankton: implications for food web ecology. Mar Biol 161:473–480. doi: 10.1007/s00227-013-2345-y

D’Ambra I, Graham WM, Carmichael RH, Hernandez FJ (2015) Fish rely on scyphozoan hosts as a primary food source: Evidence from stable isotope analysis. Mar Biol 162:247–252. doi: 10.1007/s00227-014-2569-5

de Lafontaine Y, Leggett WC (1988) Predation by jellyfish on larval fish: an experimental evaluation employing in situ enclosures. Mar Ecol Prog Ser 42:13–20.

Dittrich B (1988) Studies on the life cycle and reproduction of the parasitic amphipod *Hyperia galba* in the North Sea. Helgolander Meeresunters 42:79–98. doi: 10.1007/BF02364205

Dodge KL, Galuardi B, Lutcavage ME (2011) Jellyfish as prey for leatherback sea turtles in the North Atlantic. Mar Biol 158:225–239. doi:10.1007/s00227-010-1556-6

Doyle TK, Hay GC, Harrod C, Houghton J (2014) Ecological and societal benefits of gelatinous zooplankton. In: Gelatinous zooplankton blooms. Springer, Netherlands, pp 105–127

Energinet. 2024. Kattegat – Technical report on fish populations and habitats. Technical Report prepared by DNV for Energinet, Fredericia, Denmark. 68 pp.

Essink K, Kleef HL, Visser W (1989) On the pelagic occurrence and dispersal of the benthic amphipod corophium volutator. J Mar Biol Assoc UK 69:11–15. doi: 10.1017/S0025315400049067

Fleming NEC, Harrod C, Griffin DC, Newton J, Houghton JDR (2014) Scyphozoan gelatinous zooplankton provide short-term reproductive habitat for hyperiid amphipods in a temperate near-shore environment. Mar Ecol Prog Ser 510:229–240. doi: 10.3354/meps10896

Fujioka K, Wada H, Okano H (1993) Torishima whale bone deep-sea animal community assemblage. J Geogr (Chigaku Zasshi) 102:507–517. doi: 10.5026/jgeography.102.507

Gannes LZ, O’Brien DM, del Rio CM (1997) Stable isotopes in animal ecology: assumptions, caveats, and a call for more laboratory experiments. Ecology 78:1271–1276. doi: 10.1890/0012-9658(1997)078[1271:SIIAEA]2.0.CO;2

Garstecki T, Wickham SA (2001) Effects of resuspension and mixing on population dynamics and trophic interactions in a model benthic microbial food web. Aquat Microb Ecol 25:281–292. doi: 10.3354/ame025281

Gasca R, Haddock SHD (2004) Associations between gelatinous zooplankton and hyperiid amphipods (Crustacea: Peracarida) in the Gulf of California. Hydrobiologia 530:529–535. doi: 10.1007/s10750-004-2657-5

Gillet P, Surugiu V, Vasile R, Metais I, Mouloud M, Simo P (2011) Preliminary data on population dynamics and genetics of *Alitta succinea* (Polychaeta: Nereididae) from the Romanian coast of the Black Sea. Ital J Zool, 78: 229–241. 10.1080/11250003.2011.593347

Grey J, Jones RI, Sleep D (2001) Seasonal changes in the importance of the source of organic matter to the diet of zooplankton in Loch Ness, as indicated by stable isotope analysis. Limnol Oceanogr 46:505–513. doi: 10.4319/lo.2001.46.3.0505

Harbison GR, Biggs DC, Madin LP (1977) The associations of amphipoda Hyperiidea with gelatinous zooplankton—II. Associations with Cnidaria, Ctenophora and Radiolaria. Deep-Sea Res 24:465–488. doi: 10.1016/0146-6291(77)90484-2

Heip C, Herman R (1979) Production of *Nereis diversicolor* O. F. Müller (Polychaeta) in a shallow brackish- water pond. Estuarine Coastal Mar Sci 8:297–305. doi: 10.1016/0302-3524(79)90047-1

Hobson KA, Alisauskas RT, Clark RG (1993) Stable-nitrogen isotope enrichment in avian tissues due to fasting and nutritional stress: Implications for isotopic analyses of diet. Ornithol Appl 95:388–394. doi: 10.2307/1369361

Howe ER, Simenstad CA (2011) Isotopic determination of food web origins in restoring and ancient estuarine wetlands of the San Francisco Bay and Delta. Estuaries Coasts 34:597–617. doi: 10.1007/s12237-011-9376-8

Ingram BA, Pitt KA, Barnes P (2017) Stable isotopes reveal a potential kleptoparasitic relationship between an ophiuroid (*Ophiocnemis marmorata*) and the semaeostome gelatinous zooplankton, A. aurita. J Plankton Res 39:138–146. doi: 10.1093/plankt/fbw088

Jarman SN, McInnes JC, Faux C, Polanowski AM, Marthick J, Deagle BE, Southwell C, Emmerson L (2013) Adélie penguin diet monitoring by DNA metabarcoding reveals regional differences and trends over time. Mar Ecol Prog Ser 491:267–284. doi:10.3354/meps10499

Javidpour J, Cipriano-Maack AN, Mittermayr A, Dierking J (2016) Temporal dietary shift in Gelatinous zooplankton revealed by stable isotope analysis. Mar Biol 163:112. doi: 10.1007/s00227-016-2892-0

Kanagaraj G, Kumar PS, Morandini AC (2008) The occurrence of *Ophiocnemis marmorata* (Echinodermata: Ophiuroidea) associated with the rhizostome medusa *Rhopilema hispidum* (Cnidaria: Scyphozoa). J Ocean Univ China 7:421–424. doi: 10.1007/s11802-008-0421-6

Keenan SW, DeBruyn JM (2019) Changes to vertebrate tissue stable isotope (δ15N) composition during decomposition. Sci Rep 9:9929. doi: 10.1038/s41598-019-46368-5

Kempster B, Zanette L, Longstaffe FJ, MacDougall-Shackleton SA, Wingfield JC, Clinchy M (2007) Do stable isotopes reflect nutritional stress? Results from a laboratory experiment on song sparrows. Oecologia 151:365–371. doi: 10.1007/s00442-006-0597-7

Kiljunen M, Grey J, Sinisalo T, Harrod C, Immonen H, Jones RI (2006) A revised model for lipid- normalizing δ13C values from aquatic organisms, with implications for isotope mixing models. J Appl Ecol 43:1213–1222. doi: 10.1111/j.1365-2664.2006.01224.x

Kristensen E (1984) Life cycle, growth and production in estuarine populations of the polychaetes *Nereis virens* and *N. diversicolor*. Ecography 7:249–250. doi: 10.1111/j.1600-0587.1984.tb01128.x

Loudon JE, Sponheimer M, Sauther ML, Cuozzo FP (2007) Intraspecific variation in hair δ13C and δ15N values of ring-tailed lemurs (*Lemur catta*) with known individual histories, behavior, and feeding ecology. Am J Phys Anthropol 133:978–985. doi: 10.1002/ajpa.20605

Lucas CH, Höhn DP, Trueman CN (2025) Insights into the feeding of gelatinous zooplankton polyps in wild and laboratory conditions: do experiments provide realistic estimates of natural functional rates? Hydrobiologia. doi: 10.1007/s10750-024-05783-0

Marliave JB, Mills CE (1993) Piggyback riding by pandalid shrimp larvae on hydromedusae. Can J Zool 71:257–263. doi: 10.1139/z93-037

Masuda R, Yamashita Y, Matsuyama M (2008) Jack mackerel *Trachurus japonicus* juveniles use gelatinous zooplankton for predator avoidance and as a prey collector. Fisheries Sci 74:276–284. doi: 10.1111/j.1444-2906.2008.01522.x

McCutchan JH, Lewis WM, Kendall C, McGrath CC (2003) Variation in trophic shift for stable isotope ratios of carbon, nitrogen, and sulfur. Oikos 102:378–390.

Miron G, Kristensen E (1993) Factors influencing the distribution of nereid polychaetes: the sulfide aspect. Mar Ecol Prog Ser 93:143–153.

Mistri, M., Fiorin, R., Rossi, R., Munari, C. & Sconfietti, R. 2025. Feeding habits of the invasive ctenophore *Mnemiopsis leidyi* in the Gulf of Trieste. Water 17: 470. 10.3390/w17040470

Miyajima Y, Masuda R, Kurihara A, Kamata R, Yamashita Y, Takeuchi T (2011) Juveniles of threadsail filefish, Stephanolepis cirrhifer, can survive and grow by feeding on moon gelatinous zooplankton A. aurita. Fish Sci 77:41–48. doi: 10.1007/s12562-010-0305-8

O’Rorke R, Lavery SD, Wang M, Gallego R, Waite AM, Beckley LE, Thompson PA, Jeffs AG (2015) Phyllosomata associated with large gelatinous zooplankton: hitching rides and stealing bites. ICES J Mar Sci 72:i124–i127. doi: 10.1093/icesjms/fsu163

Ohtsuka S, Koike K, Nakaguchi K, Yamada Y, Kondo Y, Metillo EB, Boxshall GA (2009) Symbiotic copepods associated with medusae in the Philippines, with description of a new species of Paramacrochiron (Cyclopoida: Macrochironidae). Plankton Biol Ecol 56:1–13.

Pacariz S, Björk G, Svedäng H, Interannual variability in the transport of fish eggs in the Kattegat and Öresund, ICES Journal of Marine Science, Volume 71, Issue 7, September/October 2014, Pages 1706–1716, 10.1093/icesjms/fsu044

Post DM (2002) Using stable isotopes to estimate trophic position: models, methods, and assumptions. Ecology 83:703–718. doi: 10.1890/0012-9658(2002)083[0703:USITET]2.0.CO;2

Quezada-Romegialli C, Jackson AL, Hayden B, Kahilainen KK, Lopes C, Harrod C (2018) tRophicPosition, an r package for the Bayesian estimation of trophic position from consumer stable isotope ratios. Methods Ecol Evol 9:1592–1599. doi: 10.1111/2041-210X.13009

Raffaelli D, Bell E, Weithoff G, Matsumoto A, Cruz-Motta JJ, Kershaw P, Parker R, Parry D, Jones M (2003) The ups and downs of benthic ecology: considerations of scale, heterogeneity and surveillance for benthic–pelagic coupling. J Exp Mar Biol Ecol 285–286:191–203. doi: 10.1016/S0022-0981(02)00527-0

Ram JL, Fei X, Danaher SM, Lu S, Breithaupt T, Hardege JD (2008) Finding females: pheromone-guided reproductive tracking behavior by male *Nereis succinea* in the marine environment. J Exp Biol 211 (5): 757–765. doi: 10.1242/jeb.012773

Riascos JM, Docmac F, Reddin C, Harrod C (2015) Trophic relationships between the large scyphomedusa *Chrysaora plocamia* and the parasitic amphipod *Hyperia curticephala*. Mar Biol 162:1841–1848. doi: 10.1007/s00227-015-2716-7

Riisgård HU, Bondo Christensen P, Olesen NJ, Petersen JK, Møller MM, Andersen P (1995) Biological structure in a shallow cove (Kertinge Nor, Denmark) — Control by benthic nutrient fluxes and suspension-feeding ascidians and gelatinous zooplankton. Ophelia 41:329–344. doi: 10.1080/00785236.1995.10422051.

Riisgård HU, Barth-Jensen C, Madsen CV (2010) High abundance of the jellyfish *Aurelia aurita* excludes the invasive ctenophore *Mnemiopsis leidyi* to establish in a shallow cove (Kertinge Nor, Denmark). Aquat Invasions 5:347–356..

Sal Moyano MP, Schiariti A, Giberto DA, Diaz Briz L, Gavio MA, Mianzan HW (2012) The symbiotic relationship between *Lychnorhiza lucerna* (Scyphozoa, Rhizostomeae) and *Libinia spinosa* (Decapoda, Epialtidae) in the Río de la Plata (Argentina–Uruguay). Mar Biol 159:1933–1941. doi: 10.1007/s00227-012-1980-z

Schaub J, McLaskey AK, Forster I, Hunt BPV (2021) Experimentally derived estimates of turnover and modification for stable isotopes and fatty acids in scyphozoan gelatinous zooplankton. J Exp Mar Biol Ecol 545:151631. doi: 10.1016/j.jembe.2021.151631

Smith CR (1992) Whale falls: chemosynthesis on the deep seafloor. Oceanus 35:74–79.

Smith CR, Kukert H, Wheatcroft RA, Jumars PA, Deming JW (1989) Vent fauna on whale remains. Nature 341:27–28. doi: 10.1038/341027a0

Stock BC and Semmens BX. 2016. MixSIAR GUI User Manual. Version 3.1. https://github.com/brianstock/MixSIAR. doi:10.5281/zenodo.1209993.

Sullivan LJ, Gifford DJ (2004) Diet of the larval ctenophore *M. leidyi* A. Agassiz (Ctenophora, Lobata). J Plankton Res 26:417–431. doi: 10.1093/plankt/fbh033

Tenore KR, Gopalan UK (1974) Feeding efficiencies of the polychaete *Nereis virens* cultured on hard-clam tissue and oyster detritus. J Fish Res Bd Can 31:1675–1678. doi: 10.1139/f74-214

Tilves U, Fuentes VL, Milisenda G, Parrish CC, Vizzini S, Sabatés A (2018) Trophic interactions of the gelatinous zooplankton *Pelagia noctiluca* in the NW Mediterranean: evidence from stable isotope signatures and fatty acid composition. Mar Ecol Prog Ser 591:101–116. doi: 10.3354/meps12332

Towanda T, Thuesen EV (2006) Ectosymbiotic behavior of *Cancer gracilis* and its trophic relationships with its host *Phacellophora camtschatica* and the parasitoid *Hyperia medusarum*. Mar Ecol Prog Ser 315:221–236. doi: 10.3354/meps315221

Utne-Palm AC, Salvanes AGV, Currie B, Kaartvedt S, Nilsson GE, Braithwaite VA, Stecyk JAW, Hundt M, van der Bank M, Flynn B, Sandvik GK, Klevjer TA, Sweetman AK, Brüchert V, Pittman K, Peard KR, Lunde IG, Strandabø RAU, Gibbons MJ (2010) Trophic structure and community stability in an overfished ecosystem. Science 329:333–336. doi: 10.1126/science.1190708

Vervoort M, Gazave E (2022) Studying annelida regeneration using *Platynereis dumerilii*. In: Blanchoud S, Galliot B (eds) Whole-body regeneration. Methods Mol Biol, vol 2450. Humana, New York, NY. 10.1007/978-1-0716-2172-1_11

Weil J, Duguid WDP, Juanes F (2019) A hyperiid amphipod acts as a trophic link between a scyphozoan medusa and juvenile Chinook Salmon. Estuar Coast Shelf Sci 223:18–24. doi: 10.1016/j.ecss.2019.01.025

Wells GP, Dales RP (1951) Spontaneous activity patterns in animal behaviour: the irrigation of the burrow in the polychaetes *Chaetopterus variopedatus* Renier and *Nereis diversicolor* O. F. Müller. J Mar Biol Assoc UK 29:661–680. doi: 10.1017/S0025315400052851

