## Supplementary data for "Polychaete capture by native jellyfish and invasive ctenophore reveals a novel benthic–pelagic trophic link"

### SUPPORTING INFORMATION

**Table S1.** Number of individuals sampled at each site and season, categorized by the presence or absence of Nereididae. “Free” refers to individuals found without association with other species, while “Assoc.” refers specifically to interactions between gelatinous zooplankton and Nereididae polychaetes—either gelatinous zooplankton containing Nereididae within their tissues or Nereididae found within gelatinous zooplankton tissues. Other potential associations with taxa such as amphipods are not included in this category. Species abbreviations: AA = *A. aurita*, ML = M. *leidyi*.

| Samples/Species | Summer | | Autumn | | Total |
|---|---|---|---|---|---|
| | Site 1 | Site 2 | Site 1 | Site 2 | |
| AA free | 54 | 1 | 59 | 0 | 114 |
| AA assoc. | 33 | 0 | 5 | 0 | 38 |
| ML free | 9 | 22 | 8 | 28 | 67 |
| ML assoc. | 3 | 0 | 0 | 0 | 3 |
| Nereididae free | 5 | 0 | 0 | 0 | 5 |
| Nereididae assoc. AA | 38 | 0 | 7 | 0 | 45 |
| Nereididae assoc. ML | 3 | 0 | 0 | 0 | 3 |
| Seston | 4 | 3 | 12 | 12 | 31 |
| Zooplankton | 9 | 8 | 8 | 6 | 31 |

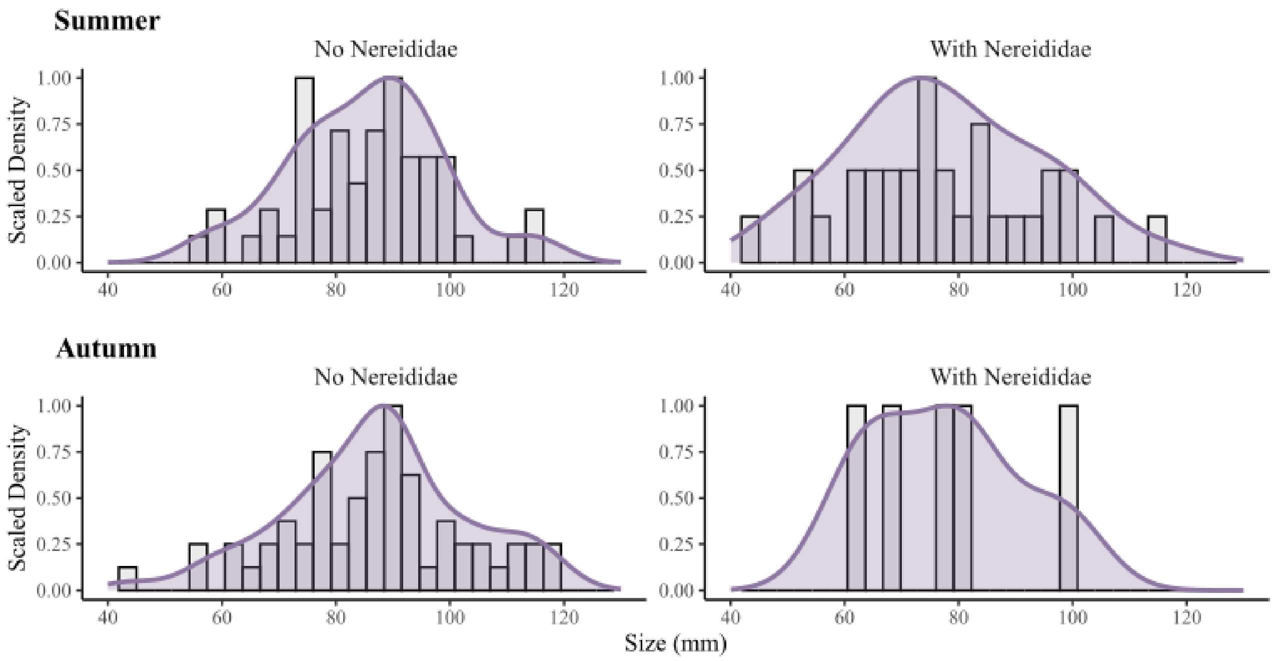


**Figure S1.** Size class density of *A. aurita* without Nereididae parasite (left) and with Nereididae found inside the gelatinous zooplankton tissue (right). Bars represent the observed frequency of medusa sizes grouped into size classes, while the overlaid smoothed curve represents the kernel density estimate, illustrating the continuous distribution pattern of sizes.

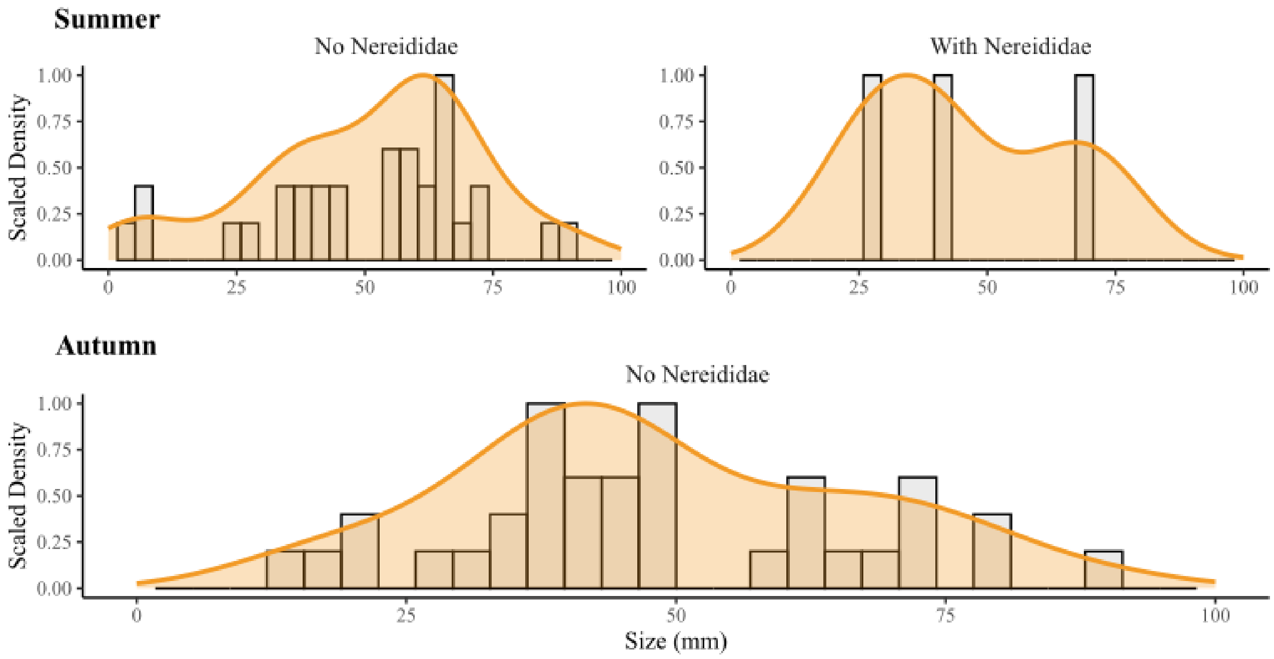


**Figure S2.** Size class density of *M. leidyi* without Nereididae parasite (left) and with Nereididae found inside the gelatinous zooplankton tissue (right). Bars represent the observed frequency of medusa sizes grouped into size classes, while the overlaid smoothed curve represents the kernel density estimate, illustrating the continuous distribution pattern of sizes.

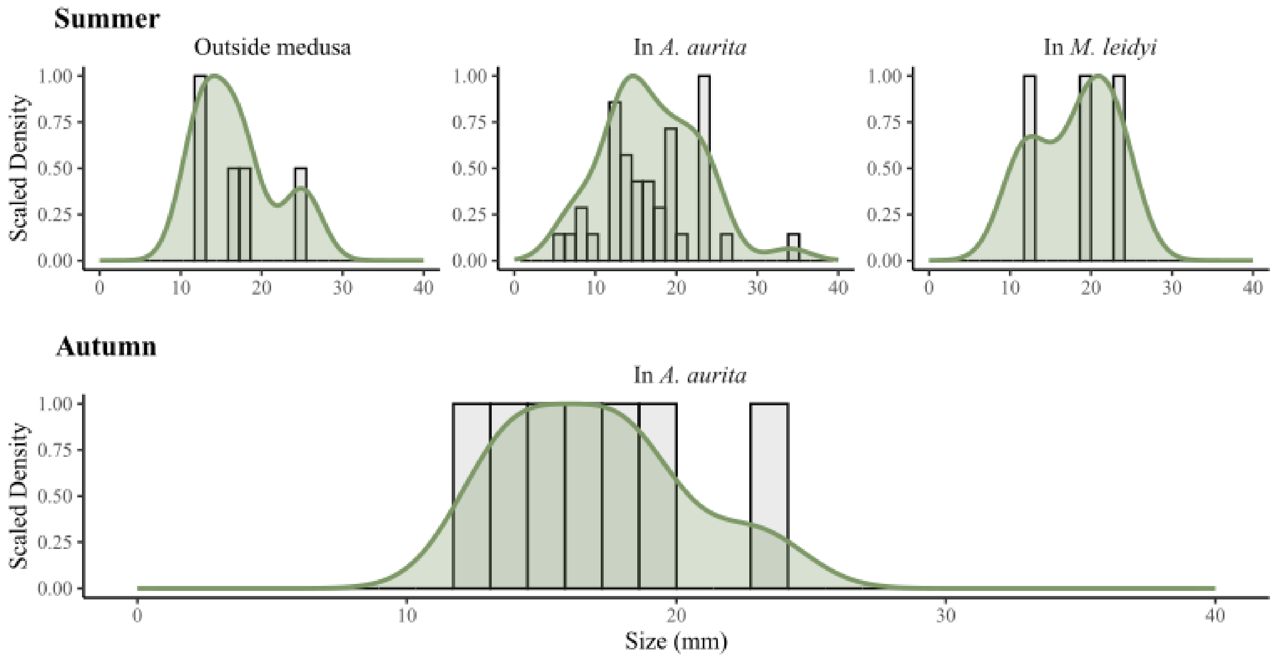


**Figure S3.** Size class density of Nereididae polychaetes outside gelatinous zooplankton, within *A.aurita* and *M. leidyi* in Summer and in Autmn. Number of samples of each depended on availbility and unique observation in the field. Bars represent the observed frequency of medusa sizes grouped into size classes, while the overlaid smoothed curve represents the kernel density estimate, illustrating the continuous distribution pattern of sizes.

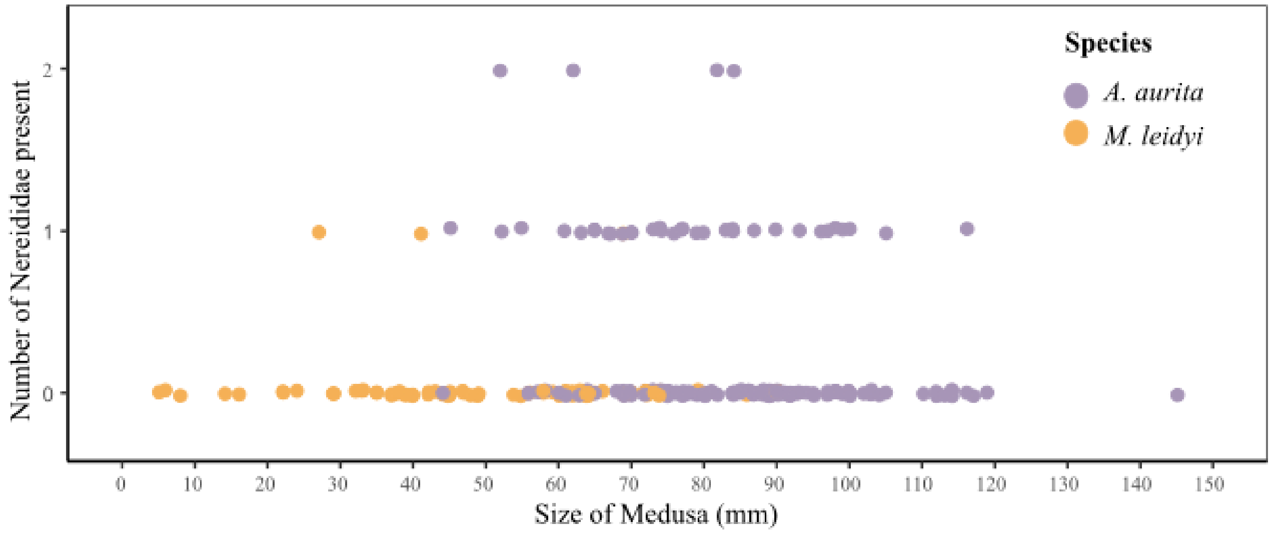

**Figure S4.** The relationship between size of medusa (*A. aurita* and *M. leidyi*) and number of Nereididae hosted by an individual gelatinous zooplankton.

**Table S2.** Mean $\delta^{13}C$ and $\delta^{15}N$ (± Standard Error) values for each gelatinous zooplankton species, Nereididae, zooplankton and seston samples during each season and at each site.

| **Season** | **Species** | **Interactions** | **Site** | **13Cmn** | **13Csd** | **13Cse** | **15Nmn** | **15Nsd** | **15Nse** |
|---|---|---|---|---|---|---|---|---|---|
| Autum | *A.aurita* | No | 1 | -18.37 | 0.94 | 0.12 | 8.45 | 1.26 | 0.16 |
| Autum | *A.aurita* | Parasites | 1 | -18.65 | 1.54 | 0.69 | 8.85 | 0.69 | 0.31 |
| Autum | *M.leidyi* | No | 1 | -20.28 | 1.60 | 0.56 | 9.87 | 0.75 | 0.27 |
| Autum | *M.leidyi* | No | 2 | -20.59 | 1.48 | 0.28 | 9.79 | 0.89 | 0.17 |
| Autum | Nereididaeae | Parasites | 1 | -14.38 | 2.24 | 0.85 | 10.64 | 2.44 | 0.92 |
| Autum | Seston | No | 1 | -22.72 | 1.99 | 0.57 | 5.45 | 1.03 | 0.30 |
| Autum | Seston | No | 2 | -22.55 | 2.03 | 0.59 | 5.90 | 0.89 | 0.26 |
| Autum | Zooplankton | No | 1 | -19.71 | 1.09 | 0.39 | 6.50 | 1.45 | 0.51 |
| Autum | Zooplankton | No | 2 | -21.06 | 1.74 | 0.71 | 6.46 | 1.20 | 0.49 |
| Summer | *A.aurita* | No | 1 | -15.66 | 1.33 | 0.18 | 8.85 | 1.03 | 0.14 |
| Summer | *A.aurita* | No | 2 | -22.54 | NA | NA | 9.52 | NA | NA |
| Summer | *A.aurita* | Parasites | 1 | -16.14 | 1.37 | 0.24 | 9.13 | 1.09 | 0.19 |
| Summer | *M.leidyi* | No | 1 | -18.03 | 1.18 | 0.39 | 10.86 | 0.96 | 0.32 |
| Summer | *M.leidyi* | No | 2 | -20.52 | 1.46 | 0.31 | 9.80 | 0.95 | 0.20 |
| Summer | *M.leidyi* | Parasites | 1 | -17.18 | 0.35 | 0.20 | 10.65 | 0.50 | 0.29 |
| Summer | Nereididaeae | No | 1 | -15.12 | 4.18 | 1.87 | 7.59 | 1.35 | 0.60 |
| Summer | Nereididaeae | Parasites | 1 | -15.47 | 3.10 | 0.48 | 8.75 | 2.17 | 0.34 |
| Summer | Seston | No | 1 | -23.82 | 2.59 | 1.30 | 6.38 | 0.91 | 0.45 |
| Summer | Seston | No | 2 | -23.70 | 1.69 | 0.97 | 5.44 | 0.29 | 0.17 |
| Summer | Zooplankton | No | 1 | -21.92 | 2.22 | 0.74 | 6.80 | 1.16 | 0.39 |
| Summer | Zooplankton | No | 2 | -23.11 | 2.31 | 0.82 | 7.20 | 1.63 | 0.58 |

**Table S3. Site 1 during summer: summary statistics for quasi-GLM analyses and pairwise comparisons of stable isotopes.** The table presents *p*-values for species pairwise comparisons of $\delta^{13}C$ (lower left, below the diagonal grey box: "–") and $\delta^{15}N$ (upper right, above the diagonal grey box: "–"). Significant differences ($p < 0.05$) between species pairs are indicated by bold and underlined values. "Free" refers to individuals found without association with other species, while "Assoc." refers specifically to interactions between gelatinous zooplankton and Nereididae polychaetes—either gelatinous zooplankton containing Nereididae within their tissues or Nereididae found within gelatinous zooplankton tissues. Species abbreviations: AA = *A. aurita*, ML = M. *leidyi*.

| p-value (d13C\d15N) | AA free | AA assoc. | ML free | ML assoc. | Nereididae free | Nereididae assoc. AA | Nereididae assoc. ML | Zooplankton | Seston |
|---|---|---|---|---|---|---|---|---|---|
| **AA free** | - | 0.990 | **0.001** | 0.322 | 0.460 | 1.000 | **0.007** | **0.001** | **0.000** |
| **AA assoc.** | 0.983 | - | **0.039** | 0.578 | 0.234 | 1.000 | **0.002** | **0.000** | **0.000** |
| **ML free** | **0.010** | 0.185 | - | 1.000 | **0.001** | **0.008** | **0.000** | **0.000** | **0.000** |
| **ML assoc.** | 0.973 | 0.999 | 1.000 | - | **0.032** | 0.421 | **0.000** | **0.001** | **0.000** |
| **Nereididae free** | 0.999 | 0.968 | 0.171 | 0.920 | - | 0.375 | 0.780 | 1.000 | 0.650 |
| **Nereididae assoc. AA** | 0.986 | 0.656 | **0.002** | 0.879 | 1.000 | - | **0.005** | **0.001** | **0.000** |
| **Nereididae assoc. ML** | **0.000** | **0.000** | **0.000** | **0.001** | **0.028** | **0.001** | - | 0.896 | 1.000 |
| **Zooplankton** | **0.000** | **0.000** | **0.000** | 0.138 | **0.000** | **0.000** | **0.000** | - | 0.693 |
| **Seston** | **0.000** | **0.000** | 0.130 | 0.382 | **0.001** | **0.000** | **0.000** | 1.000 | - |

**Table S4. Site 1 during autumn: summary statistics for quasi-GLM analyses and pairwise comparisons of stable isotopes.** The table presents *p*-values for species pairwise comparisons of $\delta^{13}C$ (lower left, below the diagonal grey box: "–") and $\delta^{15}N$ (upper right, above the diagonal grey box: "–"). Significant differences ($p < 0.05$) between species pairs are indicated by bold and underlined values. "nd" denotes cases where no data were available. "Free" refers to individuals found without association with other species, while "Assoc." refers specifically to interactions between gelatinous zooplankton and Nereididae polychaetes—either gelatinous zooplankton containing Nereididae within their tissues or Nereididae found within gelatinous zooplankton tissues. Species abbreviations: AA = *A. aurita*, ML = M. *leidyi*.

| p-value (d13C\d15N) | AA free | AA assoc. | ML free | ML assoc. | Nereididae free | Nereididae assoc. AA | Nereididae assoc. ML | Zooplankton | Seston |
|---|---|---|---|---|---|---|---|---|---|
| **AA free** | - | nd | **0.009** | nd | nd | **0.000** | nd | **0.000** | **0.000** |
| **AA assoc.** | nd | - | 0.791 | nd | nd | 0.168 | nd | **0.005** | **0.000** |
| **ML free** | 0.231 | 0.840 | - | nd | nd | 0.616 | nd | **0.000** | **0.000** |
| **ML assoc.** | nd | nd | nd | - | nd | nd | nd | nd | nd |
| **Nereididae free** | nd | nd | nd | nd | - | nd | nd | nd | nd |
| **Nereididae assoc. AA** | **0.000** | **0.002** | **0.000** | nd | nd | - | nd | **0.000** | **0.000** |
| **Nereididae assoc. ML** | nd | nd | nd | nd | nd | nd | - | nd | nd |
| **Zooplankton** | 0.166 | 0.748 | 0.999 | nd | nd | **0.000** | nd | - | 0.412 |
| **Seston** | 0.915 | 0.996 | 0.766 | nd | nd | **0.000** | nd | 0.722 | - |

**Table S5. Site 2 during Summer: summary statistics for quasi-GLM analyses and pairwise comparisons of stable isotopes.** The table presents *p*-values for species pairwise comparisons of $\delta^{13}C$ (lower left, below the diagonal grey box: “–”) and $\delta^{15}N$ (upper right, above the diagonal grey box: “–”). Significant differences ($p < 0.05$) between species pairs are indicated by bold and underlined values. “Free” refers to individuals found without association with other species, while “Assoc.” refers specifically to interactions between gelatinous zooplankton and Nereididae polychaetes—either gelatinous zooplankton containing Nereididae within their tissues or Nereididae found within gelatinous zooplankton tissues. Species abbreviations: AA = *A. aurita*, ML = M. *leidyi*.

| p-value (d13C\d15N) | AA free | AA assoc. | ML free | ML assoc. | Nereididae free | Nereididae assoc. AA | Nereididae assoc. ML | Zooplankton | Seston |
|---|---|---|---|---|---|---|---|---|---|
| **AA free** | - | 0.990 | **0.001** | 0.322 | 0.460 | 1.000 | **0.007** | **0.001** | **0.000** |
| **AA assoc.** | 0.983 | - | **0.039** | 0.578 | 0.234 | 1.000 | **0.002** | **0.000** | **0.000** |
| **ML free** | **0.010** | 0.185 | - | 1.000 | **0.001** | **0.008** | **0.000** | **0.000** | **0.000** |
| **ML assoc.** | 0.973 | 0.999 | 1.000 | - | **0.032** | 0.421 | **0.000** | **0.001** | **0.000** |
| **Nereididae free** | 0.999 | 0.968 | 0.171 | 0.920 | - | 0.375 | 0.780 | 1.000 | 0.650 |
| **Nereididae assoc. AA** | 0.986 | 0.656 | **0.002** | 0.879 | 1.000 | - | **0.005** | **0.001** | **0.000** |
| **Nereididae assoc. ML** | **0.000** | **0.000** | **0.000** | **0.001** | **0.028** | **0.001** | - | 0.896 | 1.000 |
| **Zooplankton** | **0.000** | **0.000** | **0.000** | 0.138 | **0.000** | **0.000** | **0.000** | - | 0.693 |
| **Seston** | **0.000** | **0.000** | 0.130 | 0.382 | **0.001** | **0.000** | **0.000** | 1.000 | - |

**Table S6. Site 2 during Autumn: summary statistics for quasi-GLM analyses and pairwise comparisons of stable isotopes.** The table presents *p*-values for species pairwise comparisons of $\delta^{13}C$ (lower left, below the diagonal grey box: "–") and $\delta^{15}N$ (upper right, above the diagonal grey box: "–"). Significant differences ($p < 0.05$) between species pairs are indicated by bold and underlined values. "nd" denotes cases where no data were available. "Free" refers to individuals found without association with other species, while "Assoc." refers specifically to interactions between gelatinous zooplankton and Nereididae polychaetes—either gelatinous zooplankton containing Nereididae within their tissues or Nereididae found within gelatinous zooplankton tissues. Species abbreviations: AA = *A. aurita*, ML = M. *leidyi*.

| p-value (d13C\d15N) | AA free | AA assoc. | ML free | ML assoc. | Nereididae free | Nereididae assoc. AA | Nereididae assoc. ML | Zooplankton | Seston |
|---|---|---|---|---|---|---|---|---|---|
| **AA free** | - | 0.985 | **0.009** | nd | nd | **0.000** | nd | **0.000** | **0.000** |
| **AA assoc.** | 1.000 | - | 0.791 | nd | nd | 0.168 | nd | **0.005** | **0.000** |
| **ML free** | 0.231 | 0.840 | - | nd | nd | 0.616 | nd | **0.000** | **0.000** |
| **ML assoc.** | nd | nd | nd | - | nd | nd | nd | nd | nd |
| **Nereididae free** | nd | nd | nd | nd | - | nd | nd | nd | nd |
| **Nereididae assoc. AA** | **0.000** | **0.002** | **0.000** | nd | nd | - | nd | **0.000** | **0.000** |
| **Nereididae assoc. ML** | nd | nd | nd | nd | nd | nd | - | nd | nd |
| **Zooplankton** | 0.166 | 0.748 | 0.999 | nd | nd | **0.000** | nd | - | 0.412 |
| **Seston** | 0.915 | 0.996 | 0.766 | nd | nd | **0.000** | nd | 0.722 | - |
